# Distinct roles for TANGO1S domains in maintaining ER-Golgi architecture

**DOI:** 10.64898/2026.04.28.721365

**Authors:** Elizabeth A Lawrence, Lorna Hodgson, Judith Mantell, M. Esther Prada-Sanchez, Chrissy L Hammond, David J Stephens, Nicola L Stevenson

## Abstract

The endoplasmic reticulum (ER)-Golgi interface is a dynamic trafficking hub maintained in part by TANGO1, a scaffolding protein that coordinates proteins and membranes at ER exit sites (ERES). TANGO1 has two isoforms: TANGO1L, which has a lumenal SH3 domain, and TANGO1S, which lacks this domain but retains the transmembrane and cytoplasmic coiled-coil (CC), TEER, and PRD domains common to both forms. We showed previously that loss of both isoforms disrupts ER–Golgi organization more severely than TANGO1L loss alone, indicating TANGO1S is functional and can compensate. Here we dissect the role of each TANGO1 cytoplasmic domain in maintaining secretory pathway organisation by expressing TANGO1S domain-deletion mutants in TANGO1L-/S-knockout cells. We show that TANGO1 loss causes *cis*-Golgi vesiculation that cannot be rescued by TANGO1S. TANGO1S re-expression also failed to rescue COPII and Sec16 recruitment in this cell line. Meanwhile, the TEER domain is essential for the organisation of the ER, and the TEER, CC2 and PRD domain are required for a defined ERGIC. This study represents an advance towards a domain-level resolution of TANGO1S function.

**Summary statement:** In this study we perform rescue experiments in TANGO1 knockout cells to dissect the role of the TANGO1 cytoplasmic domains in maintaining the ER-ERGIC-Golgi continuum.

## Introduction

The early secretory pathway is a highly dynamic network of membranous compartments that support the biosynthesis, trafficking and post-translational modification of secreted proteins^1^. Transport begins with the co-translational import of nascent cargoes into the endoplasmic reticulum (ER) where enzymes and chaperones assist protein folding, processing and modification^2,3^. Cargoes are then recruited to specialised ribosome-free regions of the ER, known as ER exit sites (ERES), and transported to the ER-Golgi intermediate compartment (ERGIC), a tubulovesicular sorting hub for anterograde and retrograde transport between the ER and Golgi^4^. Finally, they progress to the Golgi for further modification and sorting prior to secretion. Concomitant with this anterograde transport, ER-resident proteins and membranes are retrieved from downstream compartments via retrograde transport^5^. This bidirectional membrane flux is essential for the organisation and adaptability of the secretory pathway, yet the mechanisms that maintain the ER–Golgi interface remain poorly understood^6^.

The structure of the ER-ERGIC-Golgi axis and the modes of transport between them is currently under some debate^7–9^. Proposed transport models include vesicular transport^10–13^, tubular extensions^14–16^, and direct tunnelling or contact^17–19^ and vary with respect to the predicted extent of membrane remodelling and COPII coat organisation. Common to all models is the existence of stable ERES domains - dense, dynamic clusters of vesicles and tubules that are often closely associated with the ERGIC or early Golgi^13^. ERES are generated by COPII coat recruitment^15^. Coat assembly is initiated by Sec12, a guanine nucleotide exchange factor that activates and recruits Sar1 GTPase to the ER membrane. Sar1-GTP then recruits Sec23 and Sec24 to form the inner COPII coat, and this recruits cargo and the Sec13/ Sec31 outer coat to generate budding structures. Full coat assembly enhances the GAP activity of Sec23 towards Sar1, thus completing the cycle^20^.

Although integral to ERES formation, the COPII machinery does not work alone in shaping this domain. Other proteins such as Sec16^21,22^ and TFG^23,24^ are proposed to act as scaffolds to regulate ERES stability and organisation. The *MIA* family of genes, namely TANGO1 (transport and Golgi organisation protein 1, encoded by the *MIA3* gene)^25^, cTAGE5 (cutaneous T-cell lymphoma-associated antigen 5, encoded by *MIA2*) and TALI (TANGO1-like, also encoded by *MIA2*)^14,26–31^ are also required for COPII regulation, membrane transport, ERES organisation and stability, and cargo recruitment. The role of TANGO1, has come under particular scrutiny recently for its role in the efficient export of unusually large or high-volume cargoes from the ER^18,27,31–38^ as well as general secretion^28,39^.

TANGO1^40^ is present as both a long (TANGO1L, 1907 amino acids) and short (TANGO1S, 785 amino acids) isoform^27^. Both isoforms contain a proline rich domain (PRD), two coiled-coil domains (CC1 and CC2) and a tether for the ERGIC at the ER (TEER) domain (Supplementary Figure 1A). The C-terminal PRD is essential for ERES localisation and interacts directly with Sec23/Sec24^40,41^ and Sec16^21,42^. The CC1 domain is required for TANGO1 self-association and contains the TEER domain that binds the NRZ tethering complex to recruit ERGIC membranes for ERES expansion^43–45^. Meanwhile, the CC2 domain of TANGO1 heterodimerises with cTAGE5^30,41^. In addition to these cytoplasmic domains, TANGO1L has a luminal domain responsible for cargo-recognition and recruitment. This MOTH/SH3-like domain^31,40–42^ ^46^ is well documented as interacting with the collagen-specific chaperone Hsp47 to recruit collagens to ERES^32,40,43,47^, however it may have additional roles in transport^48^.

Recent studies have shown that both TANGO1 isoforms are functional and required for secretion^39,49^ . In zebrafish, tango1L can only partially compensate for loss of tango1S, with *tango1S* mutants exhibiting abnormal ECM and skeletal patterning indicative of secretion defects^49^. In cells, loss of TANGO1L alone has only mild impacts on the secretory pathway^39^ whilst loss of both isoforms results in fragmentation of the Golgi, mislocalisation of ERGIC proteins, vesicle accumulation, and extensive secretion defects^39^. This suggests that the cytoplasmic domain shared by both isoforms is essential for the maintenance of early secretory pathway structure.

In this study we build on this knowledge by dissecting the role of individual cytoplasmic domains of TANGO1S in maintaining early secretory pathway organisation.

## Methods

### Cell maintenance and passage

Telomerase immortalised human retinal pigment epithelium (RPE1) cells were cultured under standard conditions at 37°C, 5% CO2. Cells were cultured in Dulbecco’s Modified Eagle Medium (DMEM/F12) [Gibco, 11320-033] supplemented with 10% FBS [Gibco, A5256707] for a maximum of 40 passages. Cells were passaged once they reached 80% confluence by washing once in 1x phosphate buffered saline (PBS) and incubating in 0.05% trypsin-EDTA [Gibco, 25300-054] at 37°C until detachment. Mycoplasma tests were run routinely (MWG Eurofins).

### Plasmid generation

The TANGO1L–HA construct was a gift from Vivek Malhotra, Cell and Developmental Biology, Centre for Genomic Regulation, Barcelona, Spain^50^

To make the TANGO1S constructs, DNA sequences were designed in SnapGene and ordered as synthetic gBlocks from IDT. These sequences were designed to have 30 bp overlaps either side of the EcoRI site in the backbone vector, pEGFP-C1. Each sequence had individual domains of TANGO1S deleted while keeping the remaining protein sequence in-frame (full details of these sequences below). The synthetic DNA sequences were incorporated into the vector backbone using the NEB HiFi DNA Assembly Kit [New England Biolabs, E5520s] according to the manufacturer’s instructions and the resulting plasmids were transformed into NEB 5-alpha competent cells [New England Biolabs, C2987H], plated onto kanamycin bacterial selection plates, colony cultured, and DNA extracted for sequencing with the Qiagen Mini-Prep Kit [Qiagen, 27106]. Isolated plasmid DNA was sent to Macrogen Europe for Whole Plasmid Sequencing to confirm the inclusion of the correct DNA insert (the sequenced plasmid maps from the clones used in this paper can be found in Supplementary Figure 1B-G). Plasmids have been deposited in Addgene: pEGFP-TANGO1S^TEER (Addgene ID 256131), pEGFP-TANGO1S^CC1 (Addgene ID 256129), pEGFP-TANGO1S^PRD (Addgene ID: 256127), pEGFP-TANGO1S (Addgene ID: 256126).

gBlock sequences:

Black = sequence before domain deletion
Blue = sequence after domain deletion
Grey = TEER domain

### TANGO1S Full length

TCCGGACTCAGATCTCGAGCTCAAGCTTCGatggactcagtacctgccactgtgccttctatcgccgctaccccgggggacccggaacttgtgggacccttgtctgtgctctacgcagccttcatagccaagctgctggagctagttgctacattgcctgatgatgttcagcctgggcctgatttttatggactgccatggaaacctgtatttatcactgccttcttgggaattgcttcgtttgccattttcttatggagaactgtccttgttgtgaaggatagagtatatcaagtcacggaacagcaaatttctgagaagttgaagactatcatgaaagaaaatacagaacttgtacaaaaattgtcaaattatgaacagaagatcaaggaatcaaagaaacatgttcaggaaaccaggaaacaaaatatgattctctctgatgaagcaattaaatataaggataaaatcaagacacttgaaaaaaatcaggaaattctggatgacacagctaaaaatcttcgtgttatgctagaatctgagagagaacagaatgtcaagaatcaggacttgatatcagaaaacaagaaatctatagagaagttaaaggatgttatttcaatgaatgcctcagaattttcagaggttcagattgcacttaatgaagctaagcttagtgaagagaaggtgaagtctgaatgccatcgggttcaagaagaaaatgctaggcttaagaagaaaaaagagcagttgcagcaggaaatcgaagactggagtaaattacatgctgagctcagtgagcaaatcaaatcatttgagaagtctcagaaagatttggaagtagctcttactcacaaggatgataatattaatgctttgactaactgcattacacagttgaatctgttagagtgtgaatctgaatctgagggtcaaaataaaggtggaaatgattcagatgaattagcaaatggagaagtgggaggtgaccggaatgagaagatgaaaaatcaaattaagcagatgatggatgtctctcggacacagactgcaatatcggtagttgaagaggatctaaagcttttacagcttaagctaagagcctccgtgtccactaaatgtaacctggaagaccaggtaaagaaattggaagatgaccgcaactcactacaagctgccaaagctggactggaagatgaatgcaaaaccttgaggcagaaagtggagattctgaatgagctctatcagcaaaaggagatggctttgcaaaagaaactgagtcaagaagagtatgaacggcaagaaagagagcacaggctgtcagctgcagatgaaaaggcagtttcggctgcagaggaagtaaaaacttacaagcggagaattgaagaaatggaggatgaattacagaagacagagcggtcatttaaaaaccagatcgctacccatgagaagaaagctcatgaaaactggctcaaagctcgtgctgcagaaagagctatagctgaagagaaaagggaagctgccaatttgagacacaaattattagaattaacacaaaagatggcaatgctgcaagaagaacctgtgattgtaaaaccaatgccaggaaaaccaaatacacaaaaccctccacggagaggtcctctgagccagaatggctcttttggcccatcccctgtgagtggtggagaatgctcccctccattgacagtggagccacccgtgagacctctctctgctactctcaatcgaagagatatgcctagaagtgaatttggatcagtggacgggcctctacctcatcctcgatggtcagctgaggcatctgggaaaccctctccttctgatccaggatctggtacagctaccatgatgaacagcagctcaagaggctcttcccctaccagggtactcgatgaaggcaaggttaatatggctccaaaagggccccctcctttcccaggagtccctctcatgagcacccccatgggaggccctgtaccaccacccattcgatatggaccaccacctcagctctgcggaccttttgggcctcggccacttc ctccaccctttggccctggtatgcgtccaccactaggcttaagagaatttgcaccaggcgttccaccaggaagacgggacctgcctctccaccctcggggatttttacctggacacgcaccatttagacctttaggttcacttggcccaagagagtactttattcctggtacccgattaccacccccaacccatggtccccaggaatacccaccaccacctgctgtaagagacttactgccgtcaggctctagagatgagcctccacctgcctctcagagcactagccaggactgttcacaggctttaaaacagagcccaAATTCTGCAGTCGACGGTACCGC GGGCCCG

## TANGO1S^CC1

TCCGGACTCAGATCTCGAGCTCAAGCTTCGatggactcagtacctgccactgtgccttctatcgccgctaccccgggggacccggaacttgtgggacccttgtctgtgctctacgcagccttcatagccaagctgctggagctagttgctacattgcctgatgatgttcagcctgggcctgatttttatggactgccatggaaacctgtatttatcactgccttcttgggaattgcttcgtttgccattttcttatggagaactgtccttgttgtgaaggatagagtatatcaagtcacggaacagcacaaggatgataatattaatgctttgactaactgcattacacagttgaatctgttagagtgtgaatctgaatctgagggtcaaaataaaggtggaaatgattcagatgaattagcaaatggagaagtgggaggtgaccggaatgagaagatgaaaaatcaaattaagcagatgatggatgtctctcggacacagactgcaatatcggtagttgaagaggatctaaagcttttacagcttaagctaagagcctccgtgtccactaaatgtaacctggaagaccaggtaaagaaattggaagatgaccgcaactcactacaagctgccaaagctggactggaagatgaatgcaaaaccttgaggcagaaagtggagattctgaatgagctctatcagcaaaaggagatggctttgcaaaagaaactgagtcaagaagagtatgaacggcaagaaagagagcacaggctgtcagctgcagatgaaaaggcagtttcggctgcagaggaagtaaaaacttacaagcggagaattgaagaaatggaggatgaattacagaagacagagcggtcatttaaaaaccagatcgctacccatgagaagaaagctcatgaaaactggctcaaagctcgtgctgcagaaagagctatagctgaagagaaaagggaagctgccaatttgagacacaaattattagaattaacacaaaagatggcaatgctgcaagaagaacctgtgattgtaaaaccaatgccaggaaaaccaaatacacaaaaccctccacggagaggtcctctgagccagaatggctcttttggcccatcccctgtgagtggtggagaatgctcccctccattgacagtggagccacccgtgagacctctctctgctactctcaatcgaagagatatgcctagaagtgaatttggatcagtggacgggcctctacctcatcctcgatggtcagctgaggcatctgggaaaccctctccttctgatccaggatctggtacagctaccatgatgaacagcagctcaagaggctcttcccctaccagggtactcgatgaaggcaaggttaatatggctccaaaagggccccctcctttcccaggagtccctctcatgagcacccccatgggaggccctgtaccaccacccattcgatatggaccaccacctcagctctgcggaccttttgggcctcggccacttcctccaccctttggccctggtatgcgtccaccactaggcttaagagaatttgcaccaggcgttccaccaggaagacgggacctgcctctccaccctcggggatttttacctggacacgcaccatttagacctttaggttcacttggcccaagagagtactttattcctggtacccgattaccacccccaacccatggtccccaggaatacccaccaccacctgctgtaagagacttactgccgtcaggctctagagatgagcctccacctgcctctcagagcactagccaggactgttcacaggctttaaaacagagcccaAATTCTGCAGTCGACGGTACCGCGGGCCCG

## TANGO1S^TEER

TCCGGACTCAGATCTCGAGCTCAAGCTTCGatggactcagtacctgccactgtgccttctatcgccgctaccccgggggacccggaacttgtgggacccttgtctgtgctctacgcagccttcatagccaagctgctggagctagttgctacattgcctgatgatgttcagcctgggcctgatttttatggactgccatggaaacctgtatttatcactgccttcttgggaattgcttcgtttgccattttcttatggagaactgtccttgttgtgaaggatagagtatatcaagtcacggaacagcaaatttctgagaagttgaagactatcatgaaagaaaatacagaacttgtacaaaaattgtcaaattatgaacagaagatcaaggaatcaaagaaacatgttcaggaaaccaggaaacagaatgtcaagaatcaggacttgatatcagaaaacaagaaatctatagagaagttaaaggatgttatttcaatgaatgcctcagaattttcagaggttcagattgcacttaatgaagctaagcttagtgaagagaaggtgaagtctgaatgccatcgggttcaagaagaaaatgctaggcttaagaagaaaaaagagcagttgcagcaggaaatcgaagactggagtaaattacatgctgagctcagtgagcaaatcaaatcatttgagaagtctcagaaagatttggaagtagctcttactcacaaggatgataatattaatgctttgactaactgcattacacagttgaatctgttagagtgtgaatctgaatctgagggtcaaaataaaggtggaaatgattcagatgaattagcaaatggagaagtgggaggtgaccggaatgagaagatgaaaaatcaaattaagcagatgatggatgtctctcggacacagactgcaatatcggtagttgaagaggatctaaagcttttacagcttaagctaagagcctccgtgtccactaaatgtaacctggaagaccaggtaaagaaattggaagatgaccgcaactcactacaagctgccaaagctggactggaagatgaatgcaaaaccttgaggcagaaagtggagattctgaatgagctctatcagcaaaaggagatggctttgcaaaagaaactgagtcaagaagagtatgaacggcaagaaagagagcacaggctgtcagctgcagatgaaaaggcagtttcggctgcagaggaagtaaaaacttacaagcggagaattgaagaaatggaggatgaattacagaagacagagcggtcatttaaaaaccagatcgctacccatgagaagaaagctcatgaaaactggctcaaagctcgtgctgcagaaagagctatagctgaagagaaaagggaagctgccaatttgagacacaaattattagaattaacacaaaagatggcaatgctgcaagaagaacctgtgattgtaaaaccaatgccaggaaaaccaaatacacaaaaccctccacggagaggtcctctgagccagaatggctcttttggcccatcccctgtgagtggtggagaatgctcccctccattgacagtggagccacccgtgagacctctctctgctactctcaatcgaagagatatgcctagaagtgaatttggatcagtggacgggcctctacctcatcctcgatggtcagctgaggcatctgggaaaccctctccttctgatccaggatctggtacagctaccatgatgaacagcagctcaagaggctcttcccctaccagggtactcgatgaaggcaaggttaatatggctccaaaagggccccctcctttcccaggagtccctctcatgagcacccccatgggaggccctgtaccaccacccattcgatatggaccaccacctcagctctgcggaccttttgggcctcggccacttcctccaccctttggccctggtatgcgtccaccactaggcttaagagaatttgcaccaggcgttccaccaggaagacgggacctgcctctccaccctcggggatttttacctggacacgcaccatttagacctttaggttcacttggcccaagagagtactttattcctggtacccgattaccacccccaacccatggtccccaggaatacccaccaccacctgctgtaagagacttactgccgtcaggctctagagatgagcctccacctgcctctcagagcactagccaggactgttcacaggctttaaaacagagcccaAATTCTGCAGTCGACGGTACCGCGGGCCCG

## TANGO1S^CC1 with TEER

TCCGGACTCAGATCTCGAGCTCAAGCTTCGatggactcagtacctgccactgtgccttctatcgccgctaccccgggggacccggaacttgtgggacccttgtctgtgctctacgcagccttcatagccaagctgctggagctagttgctacattgcctgatgatgttcagcctgggcctgatttttatggactgccatggaaacctgtatttatcactgccttcttgggaattgcttcgtttgccattttcttatgga gaactgtccttgttgtgaaggatagagtatatcaagtcacggaacagcaaaatatgattctctctgatgaagcaattaaatataaggataaaatcaagacacttgaaaaaaatcaggaaattctggatgacacagctaaaaatcttcgtgttatgctagaatctgagagagaacacaaggatgataatattaatgctttgactaactgcattacacagttgaatctgttagagtgtgaatctgaatctgagggtcaaaataaaggtggaaatgattcagatgaattagcaaatggagaagtgggaggtgaccggaatgagaagatgaaaaatcaaattaagcagatgatggatgtctctcggacacagactgcaatatcggtagttgaagaggatctaaagcttttacagcttaagctaagagcctccgtgtccactaaatgtaacctggaagaccaggtaaagaaattggaagatgaccgcaactcactacaagctgccaaagctggactggaagatgaatgcaaaaccttgaggcagaaagtggagattctgaatgagctctatcagcaaaaggagatggctttgcaaaagaaactgagtcaagaagagtatgaacggcaagaaagagagcacaggctgtcagctgcagatgaaaaggcagtttcggctgcagaggaagtaaaaacttacaagcggagaattgaagaaatggaggatgaattacagaagacagagcggtcatttaaaaaccagatcgctacccatgagaagaaagctcatgaaaactggctcaaagctcgtgctgcagaaagagctatagctgaagagaaaagggaagctgccaatttgagacacaaattattagaattaacacaaaagatggcaatgctgcaagaagaacctgtgattgtaaaaccaatgccaggaaaaccaaatacacaaaaccctccacggagaggtcctctgagccagaatggctcttttggcccatcccctgtgagtggtggagaatgctcccctccattgacagtggagccacccgtgagacctctctctgctactctcaatcgaagagatatgcctagaagtgaatttggatcagtggacgggcctctacctcatcctcgatggtcagctgaggcatctgggaaaccctctccttctgatccaggatctggtacagctaccatgatgaacagcagctcaagaggctcttcccctaccagggtactcgatgaaggcaaggttaatatggctccaaaagggccccctcctttcccaggagtccctctcatgagcacccccatgggaggccctgtaccaccacccattcgatatggaccaccacctcagctctgcggaccttttgggcctcggccacttcctccaccctttggccctggtatgcgtccaccactaggcttaagagaatttgcaccaggcgttccaccaggaagacgggacctgcctctccaccctcggggatttttacctggacacgcaccatttagacctttaggttcacttggcccaagagagtactttattcctggtacccgattaccacccccaacccatggtccccaggaatacccaccaccacctgctgtaagagacttactgccgtcaggctctagagatgagcctccacctgcctctcagagcactagccaggactgttcacaggctttaaaacagagcccaAATTCTGCAGTCGACGGTACCGCGGGCCCG

## TANGO1S^CC2

TCCGGACTCAGATCTCGAGCTCAAGCTTCGatggactcagtacctgccactgtgccttctatcgccgctaccccgggggacccggaacttgtgggacccttgtctgtgctctacgcagccttcatagccaagctgctggagctagttgctacattgcctgatgatgttcagcctgggcctgatttttatggactgccatggaaacctgtatttatcactgccttcttgggaattgcttcgtttgccattttcttatggagaactgtccttgttgtgaaggatagagtatatcaagtcacggaacagcaaatttctgagaagttgaagactatcatgaaagaaaatacagaacttgtacaaaaattgtcaaattatgaacagaagatcaaggaatcaaagaaacatgttcaggaaaccaggaaacaaaatatgattctctctgatgaagcaattaaatataaggataaaatcaagacacttgaaaaaaatcaggaaattctggatgacacagctaaaaatcttcgtgttatgctagaatctgagagagaacagaatgtcaagaatcaggacttgatatcagaaaacaagaaatctatagagaagttaaaggatgttatttcaatgaatgcctcagaattttcagaggttcagattgcacttaatgaagctaagcttagtgaagagaaggtgaagtctgaatgccatcgggttcaagaagaaaatgctaggcttaagaagaaaaaagagcagttgcagcaggaaatcgaagactg gagtaaattacatgctgagctcagtgagcaaatcaaatcatttgagaagtctcagaaagatttggaagtagctcttactcacaaggatgataatattaatgctttgactaactgcattacacagttgaatctgttagagtgtgaatctgaatctgagggtcaaaataaaggtggaaatgattcagatgaattagcaaatggagaagtgggaggtgaccggaatgagaagatgaaaaatcaaattaagcagatgatggatgtctctcggacacagactgcaatatcggtagttgaagaggatctaaagcttttacagcttaagctaagagcctccgtgtccactaaatgtaacctggaagaccaggtaaagaaattggaagatgaccgcaactcactacaagctgccaaagctggactggaagatgaatgcaaaaccttgaggcagaaagtggagattctgaatgagctctatcagcaaaaggagatggctttgcaaaagaaactgagtcaagaagagtatgaacggcaagaaagagagcacaggctgtcagctgcagatgaaaaggcagtttcggctgcagaggaagtaaaaacttacaagcggagaattgaagaaatggaggatgaattacagaagacagagcggtcatttaaacctgtgattgtaaaaccaatgccaggaaaaccaaatacacaaaaccctccacggagaggtcctctgagccagaatggctcttttggcccatcccctgtgagtggtggagaatgctcccctccattgacagtggagccacccgtgagacctctctctgctactctcaatcgaagagatatgcctagaagtgaatttggatcagtggacgggcctctacctcatcctcgatggtcagctgaggcatctgggaaaccctctccttctgatccaggatctggtacagctaccatgatgaacagcagctcaagaggctcttcccctaccagggtactcgatgaaggcaaggttaatatggctccaaaagggccccctcctttcccaggagtccctctcatgagcacccccatgggaggccctgtaccaccacccattcgatatggaccaccacctcagctctgcggaccttttgggcctcggccacttcctccaccctttggccctggtatgcgtccaccactaggcttaagagaatttgcaccaggcgttccaccaggaagacgggacctgcctctccaccctcggggatttttacctggacacgcaccatttagacctttaggttcacttggcccaagagagtactttattcctggtacccgattaccacccccaacccatggtccccaggaatacccaccaccacctgctgtaagagacttactgccgtcaggctctagagatgagcctccacctgcctctcagagcactagccaggactgttcacaggctttaaaacagagcccaAATTCTGCAGTCGACGGTACCGCGGGCCCG

## TANGO1S^PRD

TCCGGACTCAGATCTCGAGCTCAAGCTTCGatggactcagtacctgccactgtgccttctatcgccgctaccccgggggacccggaacttgtgggacccttgtctgtgctctacgcagccttcatagccaagctgctggagctagttgctacattgcctgatgatgttcagcctgggcctgatttttatggactgccatggaaacctgtatttatcactgccttcttgggaattgcttcgtttgccattttcttatggagaactgtccttgttgtgaaggatagagtatatcaagtcacggaacagcaaatttctgagaagttgaagactatcatgaaagaaaatacagaacttgtacaaaaattgtcaaattatgaacagaagatcaaggaatcaaagaaacatgttcaggaaaccaggaaacaaaatatgattctctctgatgaagcaattaaatataaggataaaatcaagacacttgaaaaaaatcaggaaattctggatgacacagctaaaaatcttcgtgttatgctagaatctgagagagaacagaatgtcaagaatcaggacttgatatcagaaaacaagaaatctatagagaagttaaaggatgttatttcaatgaatgcctcagaattttcagaggttcagattgcacttaatgaagctaagcttagtgaagagaaggtgaagtctgaatgccatcgggttcaagaagaaaatgctaggcttaagaagaaaaaagagcagttgcagcaggaaatcgaagactggagtaaattacatgctgagctcagtgagcaaatcaaatcatttgagaagtctcagaaagatttggaagtagctcttactcacaaggatgataatattaatgctttgactaactgcattacacagttgaatctgttagagtgtgaatctgaatctgagggtcaaaataaaggtggaaatgattcagatgaattagcaaatggagaagtgggaggtgaccggaatgagaagatgaaaaatcaaattaagcagatgatggat gtctctcggacacagactgcaatatcggtagttgaagaggatctaaagcttttacagcttaagctaagagcctccgtgtccactaaatgtaacctggaagaccaggtaaagaaattggaagatgaccgcaactcactacaagctgccaaagctggactggaagatgaatgcaaaaccttgaggcagaaagtggagattctgaatgagctctatcagcaaaaggagatggctttgcaaaagaaactgagtcaagaagagtatgaacggcaagaaagagagcacaggctgtcagctgcagatgaaaaggcagtttcggctgcagaggaagtaaaaacttacaagcggagaattgaagaaatggaggatgaattacagaagacagagcggtcatttaaaaaccagatcgctacccatgagaagaaagctcatgaaaactggctcaaagctcgtgctgcagaaagagctatagctgaagagaaaagggaagctgccaatttgagacacaaattattagaattaacacaaaagatggcaatgctgcaagaagaatctcagagcactagccaggactgttcacaggctttaaaacagagcccaAATTCTGCAGTCGACGGTACCGCGGGCCCG

## Transfection

Cells were grown to 80% confluence in DMEM + 10% FBS before a media change to OptiMEM media [Gibco, 11058-021]. Transfection mixtures were made up with Lipofectamine 2000 [Invitrogen, 11668-027] according to the manufacturer’s instructions and added to cells in a dropwise manner. After incubation in this mix for 18 hours, the cells were washed once in PBS before being fixed (for immunofluorescence) or lysed (for immunoblotting).

## Immunofluorescence

Cells to be stained were grown on 13mm (1.5 thickness) round glass coverslips [VWR, 631-0150] and gently washed once with sterile PBS and fixed in 100% MeOH for 4 minutes at 20°C. Following fixation, the cells were: washed in PBS; blocked in 3% bovine serum albumin (BSA A9647; Sigma-Aldrich) for 30 minutes; incubated in 3% BSA plus primary antibody for 1 hour at room temperature (full list of primary antibodies and their concentrations in Table 1); washed in PBS; and incubated in 3% BSA plus AlexaFluor-or Cy-conjugated secondary antibody (all used at 1:500) for 1 hour at room temperature. Coverslips were then washed with PBS and treated with 4′,6-diamidino-2-phenylindole (DAPI) [Invitrogen, D21490] for 5 minutes at room temperature prior to final PBS washes and mounting onto glass slides with Mowiol mounting medium.

**Table 1.**

| Species | Target | Company | Catalogue number | Lot number | RRID | Dilution |
| --- | --- | --- | --- | --- | --- | --- |
| **Immunogold EM** |  |  |  |  |  |  |
| Mouse | Calnexin | Santa Cruz Biotechnology | sc-23954 | A1421 | RRID:AB_626783 | 1:5 |
| Mouse | LMAN1 (ERGIC53) | Invitrogen | MA5-25345 | YH4016563 | RRID:AB_2723324 | 1:10 |
| Mouse | GM130 | BD Bioscience | BD, 610823 | 2025007 | RRID:AB_398142 | 1:10 |
| Rabbit | Sec31A | Homemade |  |  |  | 1:5 |
| **Immunofluorescence** |  |  |  |  |  |  |
| Mouse | Calnexin | Santa Cruz Biotechnology | sc-23954 | A1421 | RRID:AB_626783 | 1:100 |
| Mouse | LMAN1 (ERGIC53) | Invitrogen | MA5-25345 | YH4016563 | RRID:AB_2723324 | 1:500 |
| Mouse | GM130 | BD Bioscience | BD, 610823 | 2025007 | RRID:AB_398142 | 1:500 |
| Rabbit | KIAA0310 (Sec16A) | Bethyl | A300-648A | Unknown | RRID:AB_519338 | 1:500 |
| Rabbit | Sec24D | Homemade |  |  |  | 1:100 |
| Rabbit | Sec31A | Homemade |  |  |  | 1:100 |
| Chicken | GFP | Abcam | Ab13970 | 1018753-4 | RRID:AB_300798 | 1:500 |
| Rabbit | cTAGE5 | Proteintech | 55279-1-AP | 09000496 | RRID:AB_11183044 | 1:500 |
| Rabbit | HA | Cell Signalling | C29F4 | 10 | RRID:AB_2798368 | 1:500 |
| Mouse | HA.11 | Covance | MMS-101P | E10031BF | RRID:AB_2314672 | 1:500 |

## Electron microscopy and tomogram generation

For generation of tomograms by electron microscopy, cells were grown to 100% confluence in 10cm dishes prior to fixation in 2.5% glutaraldehyde in 0.1M cacodylate buffer for 20 minutes. Cells were then washed in 0.1M cacodylate buffer and fixed in 1% OsO4, 1.5% potassium ferro-cyanide, 0.1M cacodylate buffer for 60 minutes at room temperature. Final washes in 0.1M cacodylate buffer and water were performed before negative staining with 3% uranyl acetate for 20 minutes. Samples were then dehydrated through sequential EtOH dilutions to a final incubation in 100% EtOH. This was replaced with a 50:50 mix of propylene oxide and epon and the dish incubated for two hours under agitation, before exchanging for 100% epon and adding resin stubs. Samples were baked at 70°C for two days, stubs removed from the dish and samples trimmed before 250nm sections were cut using a diamond knife and a UC6 ultramicrotome [Leica Microsystems]. Sections were incubated with 10nm gold nanoparticles [Sigma-Aldrich #752584] to act as fiducials. Imaging was performed using a Thermo Fisher Tecnai 20 LaB6 200k kV twin lens transmission electron microscope set to collect a tilt series between −70° and +70° using a Fischione tomography holder and FEI software. Tomograms were reconstructed using IMOD software (Kremer, 1996). Alignment was computed by tracking fiducials and tomograms generated using 10 iterations with SIRT-like filter reconstruction.

## Tokuyasu sectioning and immunogold electron microscopy

Cells were prepared for immunogold electron microscopy according to the protocol published in^51^. Briefly, RPE1 cells were grown to 100% confluence in 10cm petri dishes, fixed in glutaraldehyde and pelleted in gelatine prior to block-cutting, sucrose infiltration at 4°C overnight, mounting onto metal stubs, and snap freezing in liquid nitrogen. The frozen blocks were then cryo-sectioned to give 70nm sections on plexiform coated copper grids to be stained. The sections were washed in PBS for 30 minutes at 37°C, quenched in 0.15% glycine and blocked in 3% BSA for 1 hour at room temperature prior to incubation on drops of 3% BSA plus primary antibody (antibody details in Table 1) in a humidity chamber at 4°C overnight. The grids were then washed in 3% BSA, incubated with gold nanoparticle-conjugated secondary antibodies (Aurion DAR 10nm [810.311, batch: DAR – 10610/1] or DAM 6nm [806.322, batch: DAM – 11201/1]) diluted 1:10 in BSA for 1 hour in the dark at room temperature, and washed numerous times in dH2O. Finally, the grids were counterstained in 3% uranyl acetate for 5 minutes on ice before drying overnight. Samples were imaged on a Thermo Fisher Tecnai 12 BioTwin transmission electron microscope operating at 120 kV with images recorded on a Thermo Fisher CETA 4kx4k camera.

## Image acquisition and analysis

Widefield images were acquired with an Olympus IX-71 microscope using a 60x 1.42 N.A. objective, Exfo 120 metal halide illumination with excitation, dichroic and emission filters [Semrock, Rochester, NY], and a Photometrics Coolsnap HQ2 CCD, controlled by Volocity 5.4.1 [Perkin Elmer]. Chromatic shifts in images were registration corrected using TetraSpek fluorescent beads [Thermo Fisher]. Analysis of punctae was performed in Volocity using object intensity and size as identifiers as previously described in McCaughey et al^39^ or Image J analyse particles function^52^. Golgi area was analysed from widefield images using the freehand selection tool and measure command in ImageJ. Golgi area and ER area were normalised to cell area.

## Statistical analysis

All statistical analyses were performed in GraphPad [Prism, version 10.2.0]. Sample size ‘n’ is refers to an independent experiment (cell seeding, transfection and staining). For EM: Parts of whole used to graph distribution of gold nanoparticles across organelles in C”–F” (n = 3). O vs. E analysis (Chi-square test) was performed to compare distributions between WT and TANGO1L-/S-cells for each label (observed parameter = number of nanoparticles per organelle in TANGO1L-/S-cells, expected parameter = number of nanoparticles per organelle in WT cells). For LM: D’Agostino and Pearson normality tests were performed for all datasets, followed by the appropriate statistical test. Details of specific tests performed for each experiment are listed in figure legends. In all figures, *= p £ 0.05, ** = p £ 0.01, *** = p £ 0,001, **** = p £ 0.0001.

## Results and Discussion

### TANGO1 knockout cells accumulate Golgi-derived vesicles

In our previous study^39^, we reported observing an accumulation of unidentified membranous structures throughout TANGO1L-/S-cells. Here, we set out to characterise these structures in more detail. Using electron microscopy (EM) tomography, we determined they had the spherical morphology of vesicles (Figure 1A). We then used immunogold-labelling of early secretory pathway markers to determine their membrane identity. As this approach requires Tokuyasu cryosections rather than resin-embedded samples, we first verified that the vesicles and secretory organelles remained visible using this method. Indeed, ER and Golgi membranes were identifiable in both wild-type (WT) and TANGO1L-/S-cells, with a pronounced accumulation of vesicles in the latter (Figure 1B). Sections were subsequently labelled for calnexin, Sec31A, ERGIC53, and GM130 to mark ER, COPII, ERGIC, and *cis*-Golgi membranes, respectively (Figure 1C’–F’).

**Figure 1:**
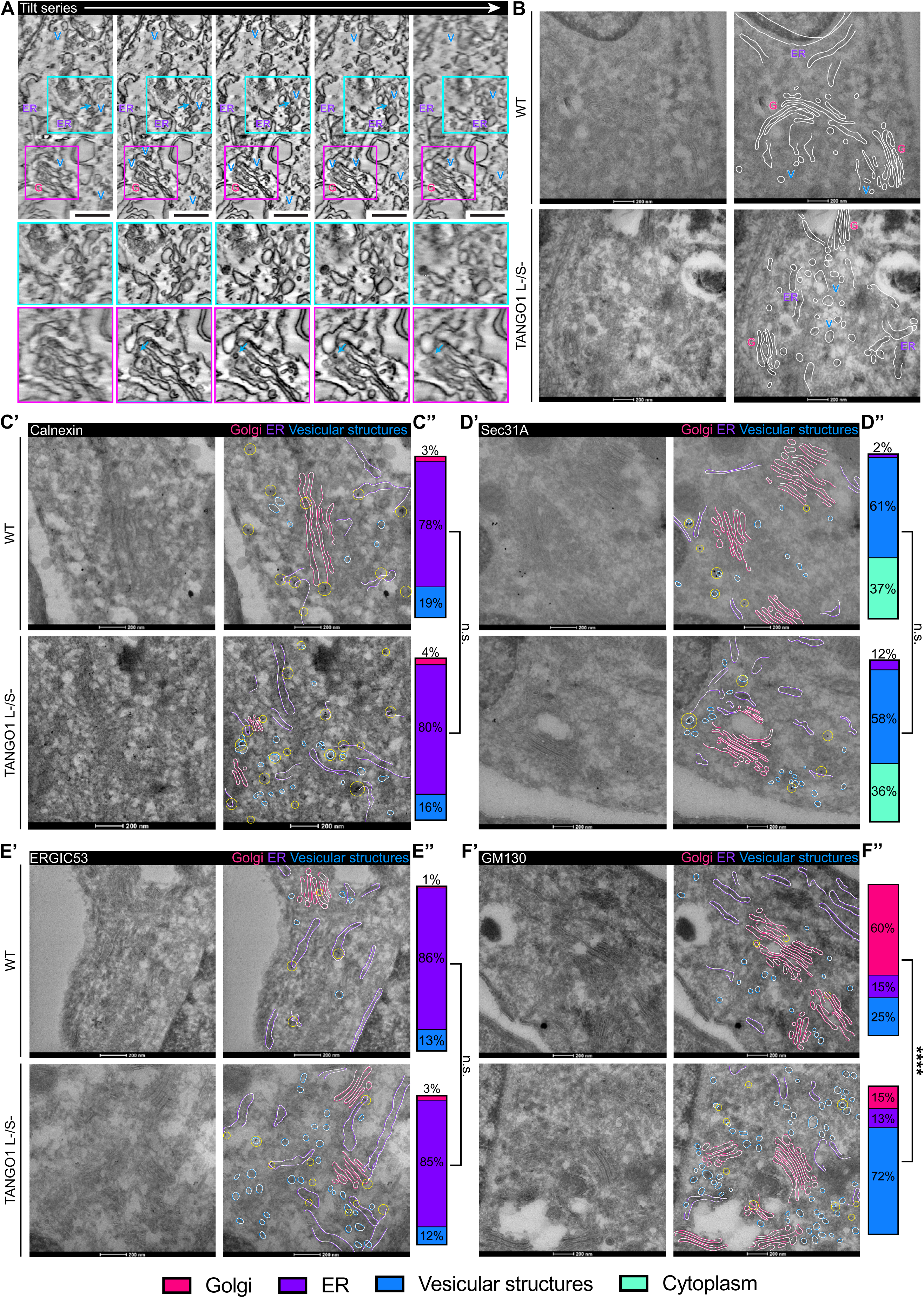
Golgi-derived vesicles accumulate in TANGO1 KO cells. **(A)** EM tomogram tilt series of TANGO1L-/S-RPE1 cells. Areas outlined by cyan and magenta boxes are enlarged in panels underneath. Arrows highlight example vesicle. Scale bar 100nm. **(B)** Electron micrographs of 70nm Tokuyasu cryosections from wildtype (WT) and TANGO1L-/S-cells. Scale bar 200nm. (A) and (B), G = Golgi, V = vesicle and ER = ER/ERGIC. **(C’–F’)** Electron micrographs of 70nm Tokuyasu cryosections from WT and TANGO1L-/S-RPE1 cells labelled with nano-Gold particles targeting Calnexin (C’), Sec31A (D’), ERGIC53 (E’), or GM130 (F’). **(C”–F”)** Distribution of gold nanoparticles across different cellular structures in WT and TANGO1L-/S-cells quantified from experiments represented in C’-F’. See methods for Statistical analysis. **(B-F)** Left-hand panel = original image, right-hand panel = annotated imaged. **(C-F)** Pink lines/bar = Golgi membranes, purple lines/bar = ER/ERGIC membranes, blue lines/bar = vesicular structures, green bars = cytoplasm, yellow circles = gold nanoparticles.

Analysis of the distribution of gold nanoparticles across early secretory pathway membranes showed that calnexin, Sec31A and ERGIC53 localisation was unchanged between WT and mutant cells (Figure 1C-E). However, labelling for GM130, the *cis-*Golgi membrane marker, was significantly altered in TANGO1L-/S-cells, with a shift in distribution from Golgi stack-associated in WT cells, to ‘vesicle-like’ structure-associated in mutant cells (Figure 1F). This agrees with previous observations that the Golgi appears fragmented in TANGO1L-/S-cells by light microscopy^39^, and suggests that TANGO1 loss causes vesiculation of the *cis*-Golgi or delivery defects for GM130-positive carriers. Note these are not mutually exclusive. Given the known roles of TANGO1 in coupling the ERES and ERGIC to sculpt membranes, we favour the former hypothesis that there is a general restructuring of the early secretory pathway. Indeed, our previous data demonstrated a collapse of the ERGIC and lack of stable well defined ERES in human cells^39^. Disruption to ER-Golgi coupling has also been reported in Drosophila^29^.

Despite disruption to the early Golgi compartment, stacked cisternal structures were still identifiable in TANGO1L-/S-cells, albeit fragmented ones, suggesting later Golgi compartments can form. This suggests some anterograde transport persists in mutant cells, as complete disruption would cause redistribution of the Golgi to the ER^53–55^. Nevertheless, stack fragmentation is consistent with broad defects in membrane flux and organisation as may be expected following ERES/ERGIC disruption.

## The luminal N-terminal domain of TANGO1L is essential for Golgi organisation

Loss of both TANGO1L and TANGO1S is more detrimental to early secretory pathway organisation than loss of a single isoform^39^. The ability of each to partially compensate for the other implies that the cytoplasmic region, common to both, is vital to preserving ER-Golgi architecture. To more specifically ascertain which cytoplasmic domains are important, we generated constructs encoding full length TANGO1S, which naturally lacks a lumenal domain, and TANGO1S with the sequence for the CC1 (^CC1), TEER (^TEER), CC1 and TEER (^CC1 and TEER), CC2 (^CC2) or PRD (^PRD) domain deleted (Supplemental Figure 1B-G). These were then expressed in the TANGO1L-/S-mutant cells to test their ability to rescue early secretory pathway organisation. Quantification of GFP intensity levels across experiments showed that TANGO1S^CC1 and TANGO1S^PRD were expressed at lower levels than other constructs (Supplemental Figure 1H). We also note that the TANGO1S deletion constructs do not show a punctate distribution, whilst high levels of TANGO1S^PRD causes aggregation.

Given our immuno-EM phenotypes, we began by looking at the Golgi. Interestingly, none of the TANGO1S constructs fully rescued Golgi structure in mutant cells. In all cases, both the overall area of the Golgi and the average fragment area was reduced in mutant cells +/- rescue when compared to WT cells (Figure 2A-C). In fact, the introduction of some domain deletion TANGO1S constructs altered Golgi morphology further, causing compaction or collapse of the membranes in some cells. Full length TANGO1S also failed to rescue Golgi fragmentation (Supplemental Figure 2A). Note whilst ^CC1 and ^PRD, appeared to reduce fragment numbers, this was through compaction rather than ribbon restoration (Supplemental Figure 2A). To test whether the long-term adaptation of the cells were preventing rescue or if the lumenal domain of TANGO1L is required, we also transfected TANGO1L into the KO lines. This did partially restore ribbon formation but not in all cells (Supplemental Figure 2B) suggesting both may be true.

**Figure 2:**
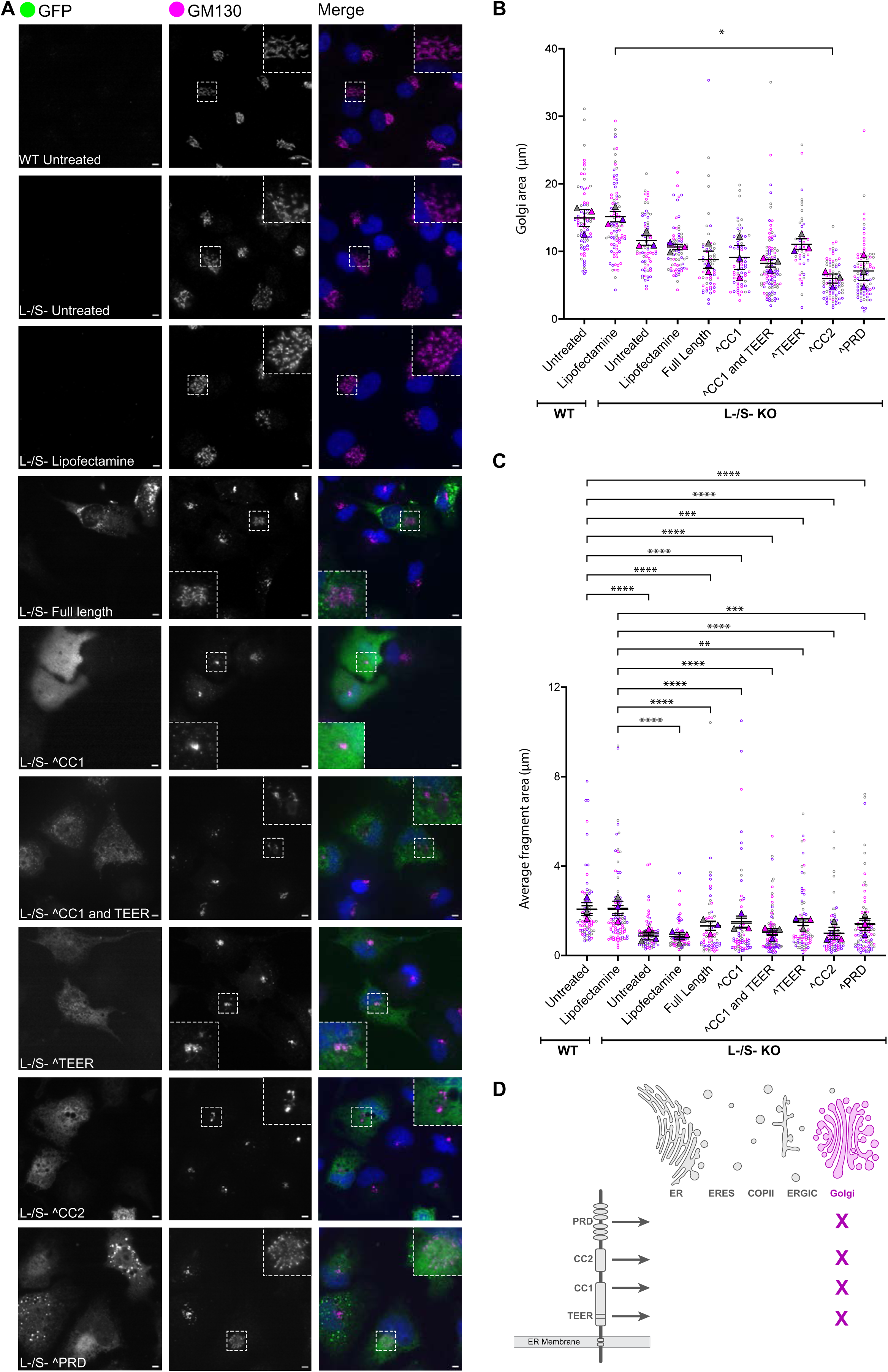
TANGO1S does not rescue Golgi phenotypes. **(A)** Images of RPE1 cells from each transfection condition immunolabelled for GFP (green) and GM130 (magenta). Inset shows zoom of area highlighted by white box. Scale bar 5µm. **(B, C)** Quantification of whole Golgi area normalised to cell area (B), and average GM130 fragment area (C) in WT, TANGO1L-/S-, and TANGO1L-/S-cells transfected with indicated constructs. Dots show individual cells colour-coded by replicate, triangles show replicate means, bars indicate overall mean and standard error of the mean, n = 3. Statistics: Kruskal-Wallis test with Dunn’s multiple comparison test. (**D)** Schematic summary of TANGO1 domains and their ability to rescue the Golgi. Crosses indicate no rescue.

The TANGO1 lumenal domain primarily binds HSP47 and collagens, raising the question of whether cargo recruitment is necessary to support membrane dynamics. The presence of cargo at ERES has been shown to influence COPII assembly^56^, and is hypothesised to sterically hinder and delay vesicle fission^57^. Both scenarios could affect transport intermediate formation and downstream compartments. Yang *et al*^48^ also recently identified an interaction between TANGO1L SH3 domain and p24 proteins in Drosophila, which act as cargo adapters^59^, regulators of ER-Golgi^60,61^ and Golgi retrograde transport^62^, and maintain Golgi architecture^63–65^. The SH3 domain may therefore have diverse roles in regulating secretion.

## The TANGO1 TEER domain is essential for ER organisation and maintenance of an ERGIC

We next investigated the role of TANGO1S in organising the ER and ERGIC membranes. In WT cells, immunolabelling for calnexin revealed a typical reticular network of well-defined tubular ER. In contrast, the ER in TANGO1L-/S-cells appeared compacted in the peri-nuclear region and calnexin staining was more diffuse, labelling thicker, denser structures (Figure 3A). Quantification of total ER area showed an increase in mutant cells compared to WT, reflecting the switch from tightly restricted narrow tubules to broader structures (Figure 3B). ER area and reticular appearance were partially rescued by all TANGO1S constructs except for those lacking the TEER domain (Figure 3A-B). The TEER domain has been shown to recruit ERGIC membranes to ERES^43,44^. Its loss may therefore impact retrograde membrane flow required to maintain the ER.

**Figure 3:**
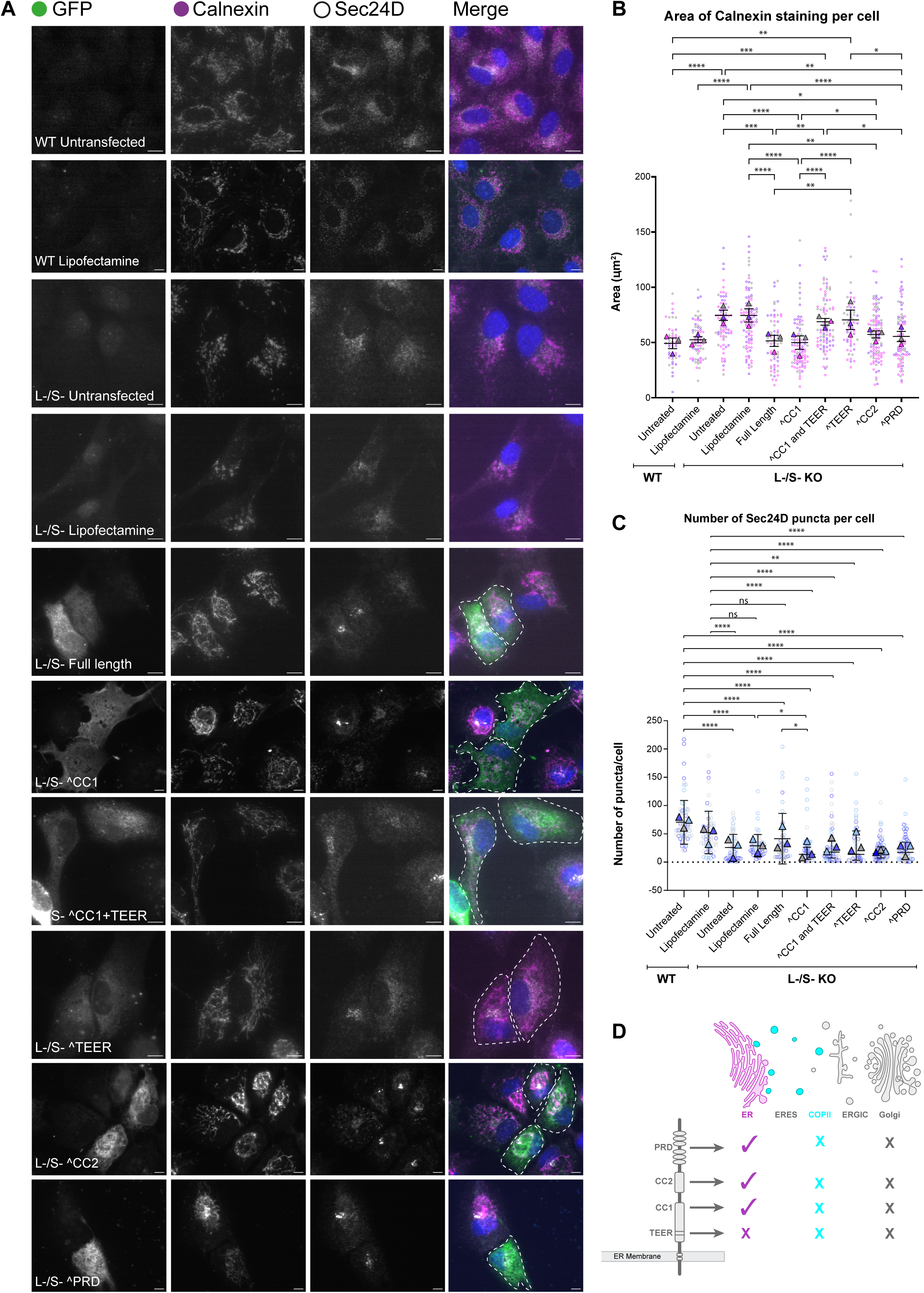
The TEER domain of TANGO1S is important for ER integrity. **(A)** Images of RPE1 cells from each transfection condition immunolabelled for GFP (green), calnexin (magenta) and Sec24D (grey). Scale bar 5µm. **(B, C)** Quantification of calnexin staining area normalised to cell area (B), and number of Sec24D punctae per cell (C) in WT, TANGO1L-/S-, and TANGO1L-/S-cells transfected as indicated. Dots show individual cells colour-coded by replicate, triangles show replicate means, bars indicate overall mean and standard error of the mean, n=3. Statistics: Kruskal-Wallis test with Dunn’s multiple comparison test. **(D)** Schematic summary of domains and their ability to rescue the ER and COPII distribution.

We next looked at the structure and organisation of the ERGIC. As previously reported, TANGO1 loss resulted in the redistribution of ERGIC marker ERGIC53 from peripheral and peri-Golgi localised puncta to ER membranes^39^ (Figure 4A). This distribution was restored upon expression of full length TANGO1S or TANGO1S^CC1 and partially restored upon TANGO1S^PRD transfection. In contrast, constructs lacking either CC2 or the TEER domain failed to rescue puncta numbers (Figure 4A-B). These domains are therefore important for the formation and/or maintenance of a distinct ERGIC.

**Figure 4:**
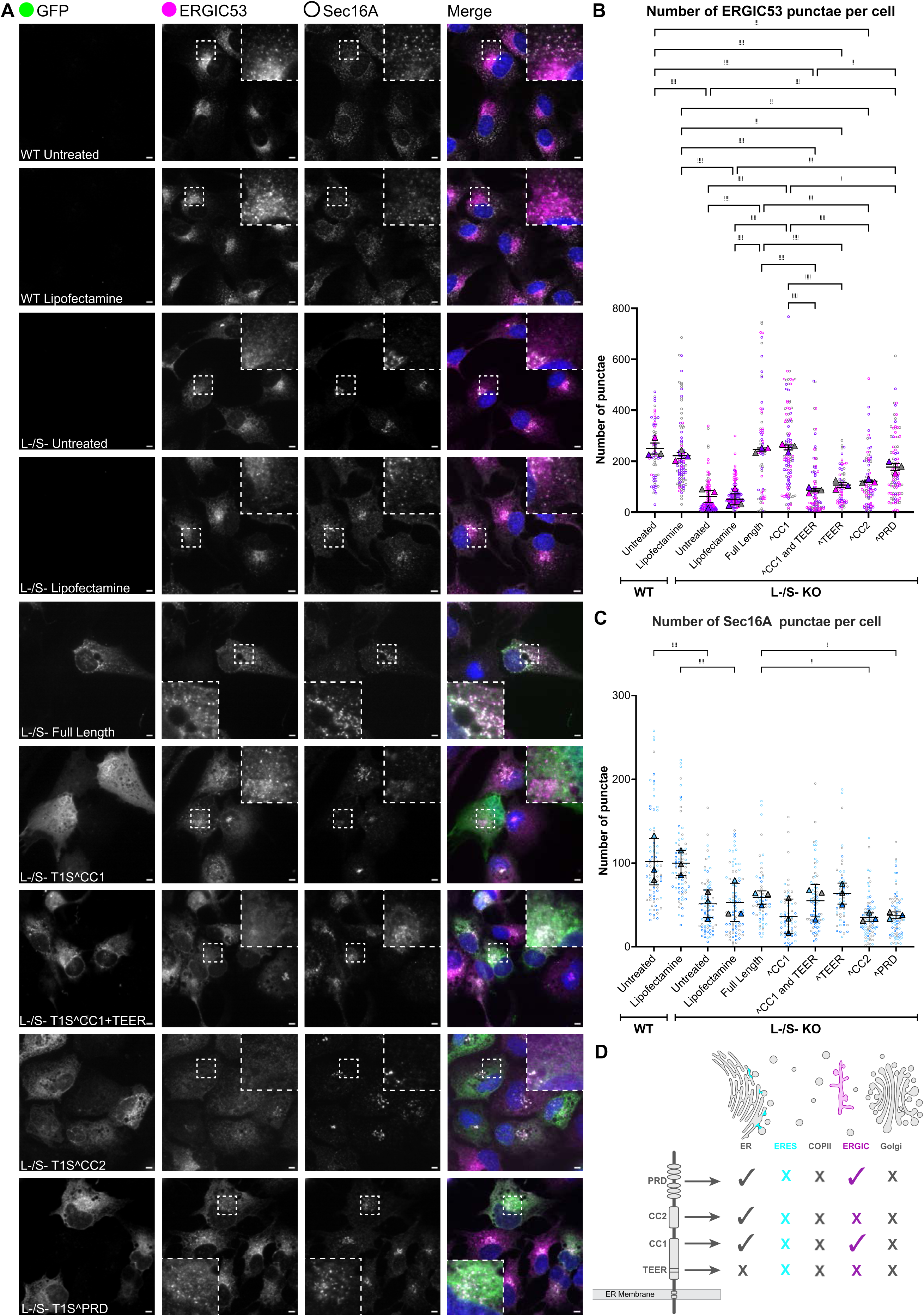
The TEER and CC2 domains of TANGO1S are important for ERGIC integrity. **(A)** Images of RPE1 cells from each transfection condition immunolabelled for GFP (green), ERGIC53 (magenta) and Sec16A (greys). Scale bar 5µm. **(B, C)** Quantification of number of ERGIC53 (B) and Sec16A punctae (C) per cell in WT, TANGO1L-/S-, and TANGO1L-/S-cells rescued with domain deletion constructs. Dots show individual cells colour-coded by replicate, triangles show replicate means, bars indicate overall mean and standard error of the mean, n=3. Statistics: One-way ANOVA with Dunn’s multiple comparison test (B), Kruskal-Wallis test with Dunn’s multiple comparison test (C). **(D)** Schematic summary of domains important for maintenance of the ERGIC and ERES.

Our current knowledge of the TEER domain is insufficient to explain its role in generating an ERGIC, however, one can speculate that this observation is still consistent with an inability to recycle ERGIC membranes and machinery effectively. Meanwhile, the CC2 domain, is best characterised for its role in oligomerising with cTAGE5^30^. Our previous investigations showed that cTAGE5 expression is significantly down-regulated in our TANGO1L-/S-line. We therefore tested whether TANGO1L or TANGO1S transfection is sufficient to restore cTAGE5 expression within the timeframe of our experiments. No cTAGE5 recovery was observed in either case, yet the ERGIC was restored (Figure 4A-B, Supplemental Figure 2C-D). It is possible that low levels of cTAGE5 persist and are sufficient for TANGO1 oligomerisation. However future investigations may need to consider cTAGE5 independent roles for the CC2 domain.

## TANGO1S does not rescue COPII recruitment or ERES organisation

We next focussed on whether TANGO1S can rescue COPII and ERES assembly by examining Sec24D and Sec16A distribution. In WT cells both proteins are punctate throughout the cell (Figure 3A and 4A), whilst in TANGO1L-/S-cells Sec24D labelling is diffuse and Sec16A redistributes to the peri-Golgi region. All TANGO1S constructs failed to rescue Sec16A or Sec24D localisation at peripheral sites (Figure 3C, 4C). TANGO1S constructs retaining the TEER domain also induced abnormal accumulation of Sec24D at the Golgi (Figure 3A). In contrast, TANGO1L did restore Sec24D puncta confirming rescue is possible in these lines (Supplemental Figure 2E). We validated these results using another COPII subunit, Sec31A, and again found that TANGO1L, but not TANGO1S could rescue COPII distribution to ERES (Supplemental Figure 2F).

These findings are consistent with the punctate versus diffuse localisation of TANGO1L and TANGO1S constructs respectively and may reflect a more general defect with TANGO1S localisation to ERES in these lines. As with the Golgi, it is also possible that the lumenal domain of TANGO1 is required for ERES organisation, however this is challenging to reconcile with other studies. For example, the PRD domain of TANGO1 has been shown to be both essential and sufficient for Sec16 recruitment^42^. Our results are consistent with the first, but not the latter. While we do not have a clear explanation for this discrepancy, we can speculate that it reflects the systems used. The Golgi accumulation of Sec24D upon TANGO1S expression could also be indicative of dominant negative effects on ERES assembly. This and likely transport defects resulting from a failure to restore ERES, may also explain the failure of TANGO1S to rescue the Golgi.

## Conclusions and study limitations

Altogether, our results underscore the pivotal role of TANGO1 in maintaining the integrity of the ER–Golgi interface, with its influence extending well beyond ERES. Our results indicate that whilst TANGO1S can rescue the ER and ERGIC, ERES and Golgi structure requires TANGO1L. There are some limitations to the study. Firstly, we are transiently over-expressing proteins into knockout lines that have undergone long-term adaptation^39^ . Our cTAGE5 data indicate this is not reversed in the time course of our experiments. The balance of machinery is therefore unlikely to reflect a ‘healthy’ state, which may impact TANGO1S behaviour. Secondly, we use the term ‘rescue’ to suggest a trend towards wildtype state only, not full restoration of membrane architecture and function. These limitations may explain why the Golgi is only partially rescued by TANGO1L. Thirdly, the deletion constructs show variable expression and localisation patterns and we may be reporting on recruitment defects. Finally, our limited understanding of the complex biology of the ER-ERGIC-Golgi continuum constrains mechanistic interpretation of our observations^8^. Nevertheless, this study represents an advance towards a domain-level resolution of TANGO1S function and offers new insights into early secretory pathway organisation.

## Acknowledgements

The authors gratefully acknowledge and thank the Wolfson Bioimaging Facility for their support and assistance in this work. We are saddened to report that Professor David Stephens died in 2024, prior to the submission of this manuscript, however we would like to express our gratitude for all his ideas, mentorship and enthusiasm.

## Author CRediT

Conceptualisation: DJS. Data curation: NLS, EAL. Formal analysis: EAL, NLS. Funding acquisition: DJS CLH NLS. Investigation: EAL JM LH EPS NLS. Project administration: DLS CLH NLS. Supervision: DJS CLH NLS. Visualisation: EAL NLS. Writing – original draft: EAL NLS. Writing – review and editing: EAL, LH, EPS, CLH, NLS.

## Funding

This research was supported by a grant from the Biotechnology and Biological Sciences Research Council (BB/V004352/1) and Medical Research Council (UKRI1419). EM was supported by BBSRC Alert 22 equipment grant (BB/X019799/1).

**Supplementary Figure 1: TANGO1 domains and constructs used in rescue experiments. (A)** Schematic representation of TANGO1L and TANGO1S protein domains and their known binding partners. **(B-G)** Maps of the pEGFP-C1 vector containing full length TANGO1S (B) or TANGO1S with CC1 and TEER (C), CC1 (D), TEER (E), CC2 (F), or PRD (G) domains missing. Schematics of the TANGO1S insert encoded shown underneath. **(H)** Quantification of GFP intensity following transfection with each construct. Dots represent individual cells, bars mean and standard deviation. N=2. Kruskal-Wallis test with Dunn’s multiple comparison test.

**Supplementary Figure 2: TANGO1L rescues the Golgi ribbon and ERES. (A)** Quantification of Golgi fragment number in TANGO1S rescue experiments (Figure 2). **(B)** Images of (i) and quantification of Golgi fragment number in (ii) cells from indicated transfection condition stained for GM130 and HA. **(C).** cTAGE5 labelling in WT, TANGO1L-/S-, and TANGO1L-/S-cells transfected with TANGO1L-HA or TANGO1S-GFP. **(D-E)** Images of (i) and quantification of puncta in (ii) cells from indicated transfection condition stained for ERGIC53 (D) or Sec24D (E). **(F)** Images of RPE1 cells from each transfection condition immunolabelled for Sec31A. **(B-F)** Images: Scale 10µm. White dash line = transfected cell. Graphs: dots = individual cells, triangles = replicate mean, bars = mean and SEM, colours indicate replicates. Statistics: Kruskal-Wallis test with Dunn’s multiple comparison test (n=3).

