## Supplementary figures and images for "Distinct roles for TANGO1S domains in maintaining ER-Golgi architecture"

### Supplemental Figure 1

#### Supplemental Figure 1

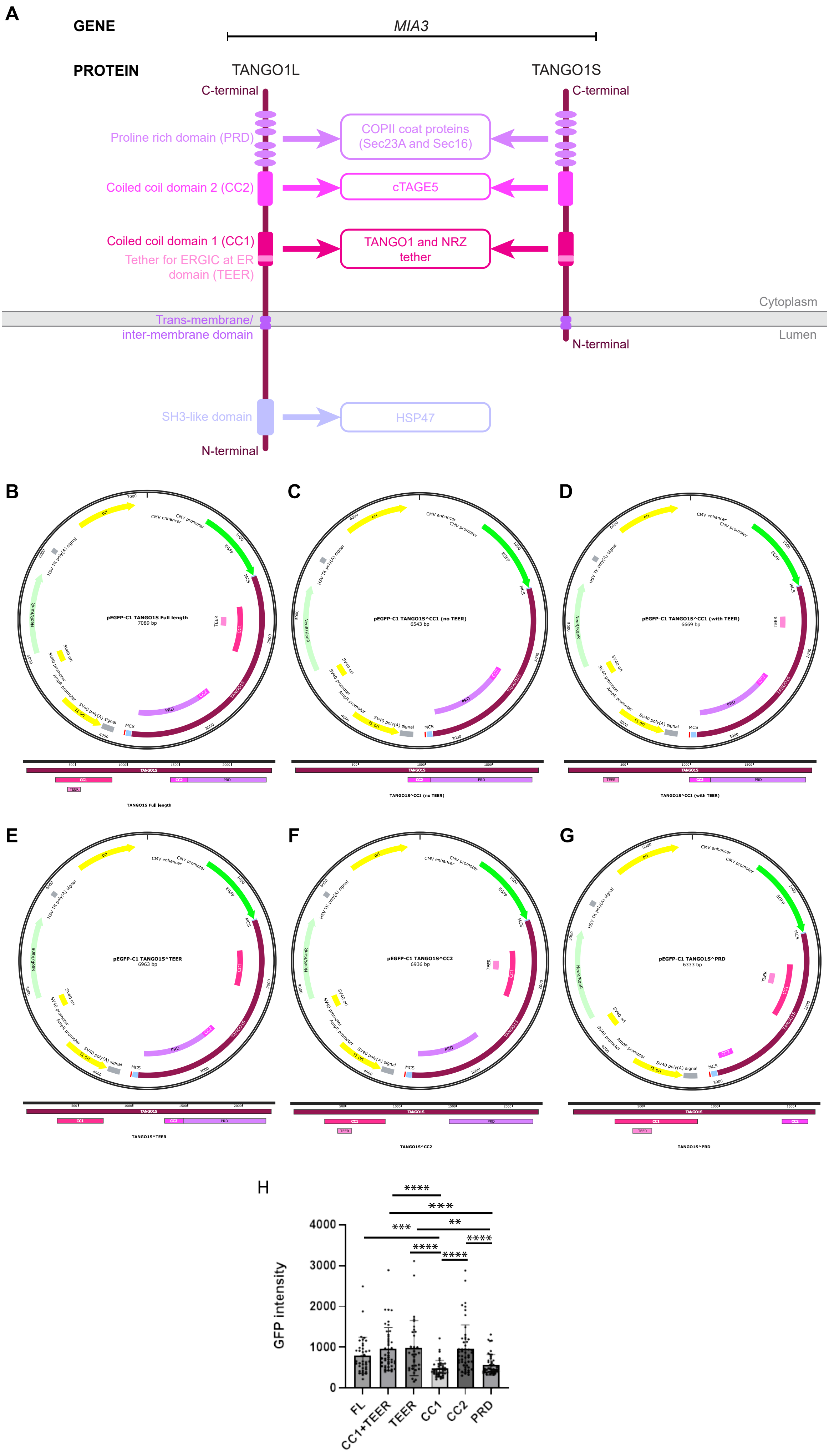

### Supplemental Figure 2

#### Supplemental Figure 2

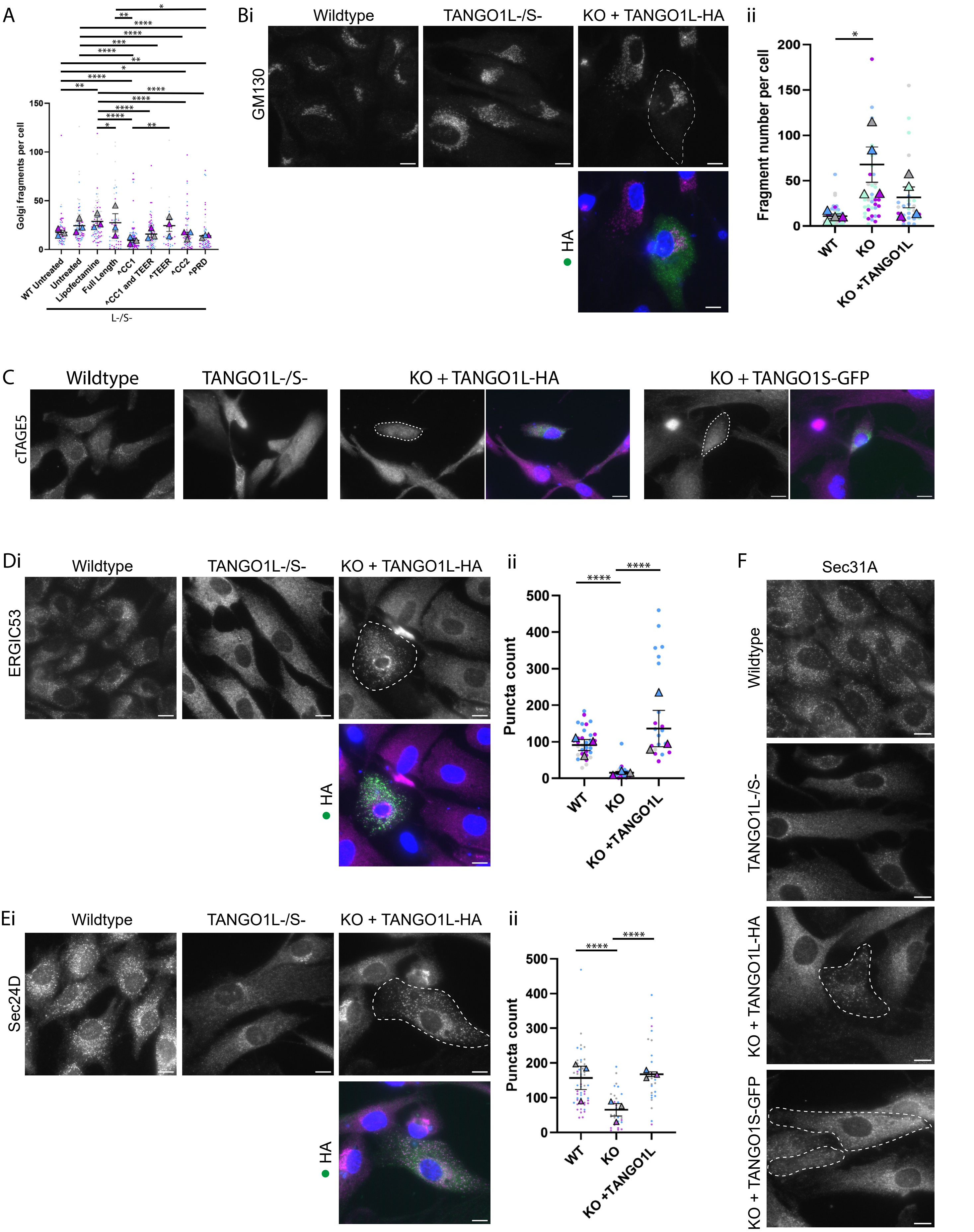
